# Efficient learning in children with rapid GABA boosting after training

**DOI:** 10.1101/2022.01.02.474022

**Authors:** Sebastian M. Frank, Markus Becker, Andrea Qi, Patricia Geiger, Ulrike I. Frank, Wilhelm M. Malloni, Yuka Sasaki, Mark W. Greenlee, Takeo Watanabe

## Abstract

It is unclear why and how children learn more efficiently than adults, although inhibitory systems, which play an important role in stabilizing learning, are immature in children. Here, we found that despite a lower baseline concentration of γ-aminobutyric acid (GABA) in early visual cortical areas in children (8 to 11 years old) than adults (18 to 35 years old), children exhibited a rapid boost of GABA immediately after visual training, whereas the concentration of GABA in adults remained unchanged after training. Moreover, behavioral experiments showed that children stabilized visual learning much faster than adults, showing rapid development of resilience to retrograde interference. These results together suggest that inhibitory systems in children’s brains are more dynamic and adapt more quickly to stabilize learning than in adults.

**One Sentence Summary:** Children learn more efficiently than adults due to faster stabilization of learning with rapid GABA boosting after training.

## Main Text

It is generally thought that school children learn more efficiently than adults. However, neuroscientifically this notion has yet to be proven. If it is true, it is unclear which aspect of learning is more efficient in children than adults. One crucial aspect of efficient learning is to learn more items within a given time. To accomplish this, it is necessary to rapidly stabilize a post-training fragile and plastic learning state and turn it into a robust memory trace that is not retrogradely interfered with by new learning. GABAergic inhibitory processing plays an important role in the stabilization of newly learned material (*1–5*). However, developmental studies have shown that the GABAergic inhibitory system is not matured yet in children (*6–9*), suggesting that children do not induce the same amount of GABAergic inhibitory processing as adults. This appears to be inconsistent with the general assumption of more efficient learning in children than adults. However, a potential problem is that GABAergic inhibitory processing in children has never been measured in the context of learning. Moreover, it remains unknown whether and, if so, how GABAergic responses occur at all in children’s brains during the stabilization of learning, that is, after task training has been terminated.

Here, we measured the concentration of GABA in early visual cortical areas of children and adults using magnetic resonance spectroscopy (MRS) before, during and after visual training. We found that the concentration of GABA was indeed lower in children than in adults until the end of training, as predicted from the immaturity of the GABAergic inhibitory system in children (*6–9*). However, to our surprise, after training, children exhibited a rapid boost in the concentration of GABA, while post-training GABA levels remained unchanged in adults. Furthermore, behavioral experiments showed that resilience to retrograde interference and therefore stabilization indeed occurred within minutes after training ended in children, whereas learning was in a fragile state in adults for at least one hour after training. The results of these experiments together suggest that, compared with adults, children exhibit a more dynamic GABAergic inhibitory system, which more rapidly adapts to stabilize learning than in adults. This rapid stabilization of learning in children enables them to learn more items within a given period of time and makes learning more efficient in children than in adults.

As mentioned above, several studies have found that GABAergic inhibitory processing is not fully matured yet in children (*6–9*), leading to the prediction that children exhibit lower baseline concentrations of GABA and overall weaker GABAergic responses than adults. To test whether this prediction is correct for GABAergic processing related to learning, we used MRS to measure the concentration of GABA in early visual cortical areas in children (n = 13) and adults (n = 14), prior to, during and after training on a visual perceptual learning (VPL) task (Fig. 1A-C). The sample size for the experiment was determined by means of a power analysis based on the results from a preliminary experiment (see *10*).

**Fig. 1.**
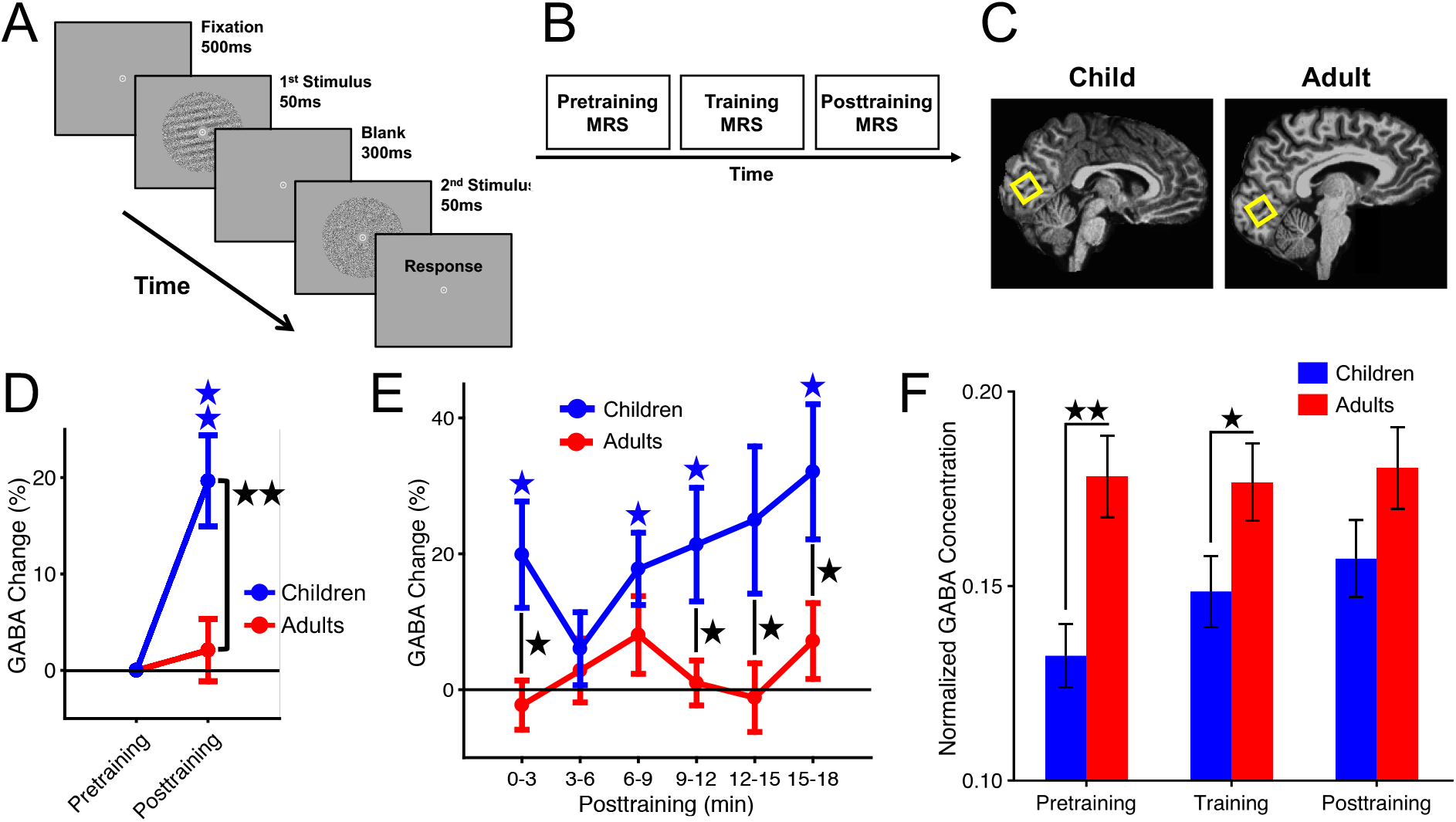
Training task and results of the MRS experiment. (**A**) Procedure of an example training trial. (**B**) Design of the experiment. (**C**) Voxel location (shown by yellow rectangle) during MRS in representative child and adult subjects. (**D**) Mean (± SEM) changes in GABA from pretraining (corresponding to zero on the y-axis) to posttraining across children (n = 13) and adults (n = 14). The blue asterisks indicate a significant increase of GABA during posttraining in children. This posttraining increase of GABA was significantly more pronounced in children than adults (indicated by black asterisks). (**E**) Same as (D) but time-resolved for every 3 min after the onset of posttraining. Statistical results after false discovery rate correction. (**F**) Mean (± SEM) concentrations of GABA normalized to a control metabolite (see *10*) across children and adults during pretraining, training and posttraining. * *p* < 0.05, ** *p* < 0.01.

We found that children exhibited a significant increase of GABA from pretraining to posttraining [one-sample *t*-test of percent change of GABA in posttraining against zero corresponding to pretraining concentrations of GABA; *t*(12) = 4.16, *p* = 0.001, *d* = 1.15], whereas GABA remained unchanged in adults [*t*(13) = 0.65, *p* = 0.53] (Fig. 1D; see Fig 1E for a time-resolved analysis of posttraining changes of GABA). This change of GABA from pretraining to posttraining was significantly different between children and adults [two-sample *t*-test; *t*(25) = 3.11, *p* = 0.005, *d* = 1.20] (Fig. 1D). For the first time, we find that the concentration of GABA in children rapidly boosted after training, whereas it did not do so in adults, although significant learning occurred in both children and adults [one-sample *t*-test of percent performance change in posttest against zero corresponding to pretest performance; children: *t*(12) = 4.12, *p* = 0.001, *d* = 1.14; adults: *t*(13) = 3.55, *p* = 0.004, *d* = 0.95]. The magnitude of learning was not significantly different between the two age groups [two-sample *t*-test; *t*(25) = 0.46, *p* = 0.65] (see *10* and fig. S1-6 for details and results of MRS control analyses).

The results (Fig. 1F) also show that, compared with adults, children exhibited significantly lower concentrations of GABA during pretraining [two-sample *t*-test; *t*(25) = −3.45, *p* = 0.002, *d* = −1.33] and training [two-sample *t*-test; *t*(24) = −2.08, *p* = 0.048, *d* = −0.81]. However, no significant difference between children and adults was found during posttraining [two-sample *t*-test; *t*(25) = −1.61, *p* = 0.12]. These results agree with the previously reported immaturity of the GABAergic inhibitory system by showing that the baseline concentration of GABA is lower in children than in adults (*6 – 9*). However, after training the concentration of GABA in children rapidly increased (see fig. S7 for time-resolved results).

Together, the results of this experiment indicate, as expected from previous studies, that the concentrations of GABA prior to and during training were lower in children than in adults (Fig. 1F and fig. S7), but that the concentration of GABA unexpectedly rapidly boosted only in children once the training was over (Fig. 1D, E and fig. S7).

Previous studies showed that a posttraining increase in the concentration of GABA is associated with the stabilization of learning (*1 – 5*). Without such stabilization, learning is in a fragile state and retrogradely interfered with by new learning (see *2, 11–15*). The rapid, posttraining boost of GABA in children found here together with the role of an increase in the concentration of GABA for the stabilization of learning lead to a prediction: Children should exhibit a more rapid stabilization of learning after training than adults. If so, VPL after training in children should be less subjected to retrograde interference, even with a short intermission before starting new learning, than VPL in adults.

To test whether this prediction is correct, we conducted a behavioral experiment, in which different groups of children (n = 14) and adults (n = 14) were trained on the orientation detection task (see Fig. 1A) using two different orientations for training. After training on the first orientation and an intermission of 10 min during which time subjects rested, they were trained on a second orientation (Fig. 2A) to test whether retrograde interference from VPL of the second orientation to VPL of the first orientation occurred. Previous studies in VPL reported that adults exhibited retrograde interference if the intermission between two different visual training sessions is 60 min or less (*2, 13*). Changes in detection thresholds from pretest, conducted immediately prior to training of the first orientation, to posttest, conducted exactly as the pretest but on a separate day after training, were calculated for each trained orientation (Fig. 2A and *10*).

**Fig. 2.**
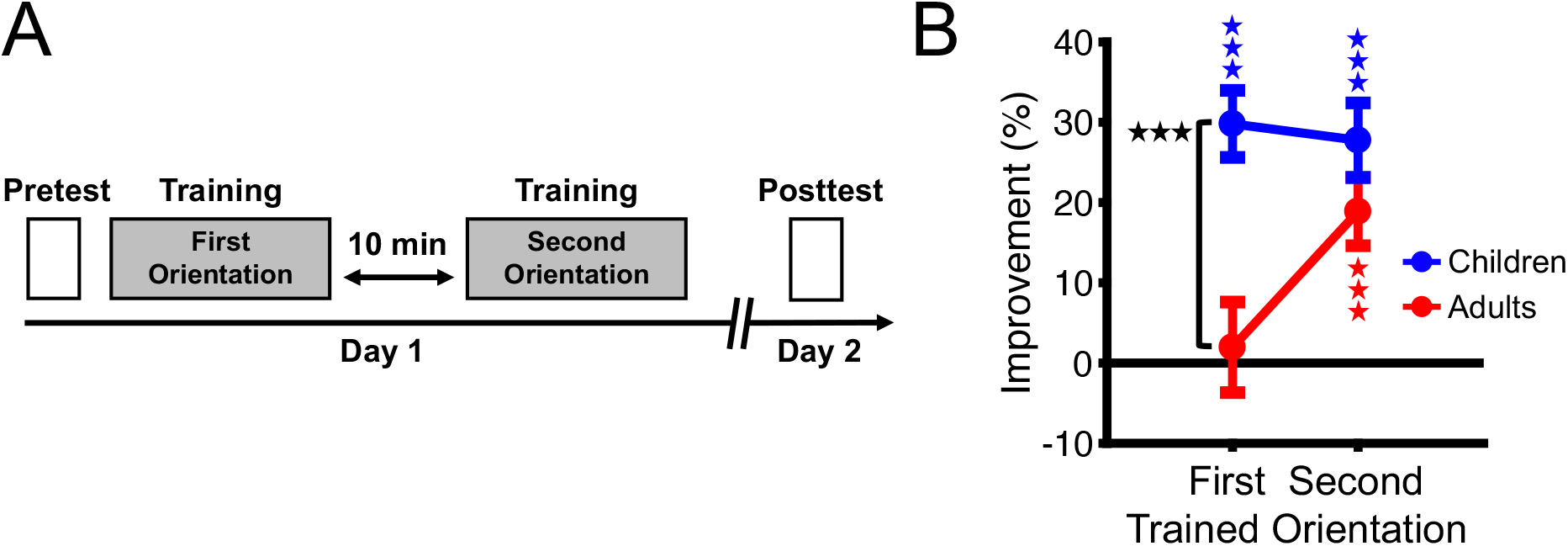
Design and results of the behavioral experiment. (**A**) Design of the experiment. (**B**) Results of the experiment. Mean (± SEM) changes in detection thresholds from pretest to posttest for each trained orientation across children (n = 14) and adults (n = 14). Zero on the y-axis represents pretest thresholds. Blue and red asterisks indicate a significant improvement in thresholds from pretest to posttest, indicative of visual perceptual learning, for children and adults, respectively. The black asterisks between children and adults for the first trained orientation indicate that children exhibited significantly greater improvements for this trained orientation than adults. *** *p* < 0.001.

Confirming our prediction, children exhibited significant VPL for each trained orientation [one-sample *t*-test of percent threshold change in posttest against zero corresponding to pretest threshold; first trained orientation: *t*(13) = 7.13, *p* < 0.001, *d* = 1.91; second trained orientation: *t*(13) = 5.96, *p* < 0.001, *d* = 1.59], indicating that no retrograde interference occurred in children (Fig. 2B). Unlike children, adults exhibited significant VPL only for the second trained orientation [*t*(13) = 4.43, *p* < 0.001, *d* = 1.18] but not for the first trained orientation [*t*(13) = 0.35, *p* = 0.73] (Fig. 2B), indicating that retrograde interference occurred in adults, similar to previously reported results for adults (*2, 13*). The difference in threshold change for the first trained orientation between children and adults was significant [two-sample *t*-test; *t*(26) = 3.97, *p* < 0.001, *d* = 1.50]. No significant difference in threshold change was found for the second trained orientation between the two age groups [*t*(26) = 1.41, *p* = 0.17]. Importantly, there were no significant differences in pretest detection threshold between children and adults for the first [*t*(26) = 1.23, *p* = 0.23] and second trained orientations [*t*(26) = −0.24, *p* = 0.81], indicating that differences in VPL for each trained orientation between children and adults did not result from any pretraining differences in thresholds between the age groups. Essentially the same pattern of results was obtained in an additional experiment using different groups of children and adults in which the duration of the intermission between training of the first and second trained orientations was 60 minutes (see fig. S8). The results of these behavioral experiments together show that children exhibited no retrograde interference with an intermission as short as 10 minutes between the first and second training, whereas adults exhibited retrograde interference with an intermission as long as 60 minutes, as reported previously (see *2, 13*).

How does an increase in the concentration of GABA occur on a cellular level? Although it is difficult to determine the cellular origin of the posttraining increase of GABA using MRS (*16–19*), it is likely that it results for the most part from an increase of synaptic GABAergic transmission and thus reflects a transient increase of neuronal inhibition in early visual cortical areas for the following reasons. First, changes in GABAergic synaptic transmissions are reflected by changes in the concentration of GABA measured by MRS (*20, 21*). Second, transient changes in GABAergic synaptic transmission by drug intake modulate learning (*22–24*). Third, training leads to transient changes in the concentrations of GABA measured by MRS (*2, 3, 5, 25, 26*).

Numerous studies indicated that children’s GABAergic inhibitory systems are not fully matured yet. However, hardly any study measured brain activation related to inhibitory processing in children (but see *27*). Moreover, no study systematically measured the concentration of GABA in children in the context of learning and compared it with adults. As a result of measuring the concentration of GABA in early visual cortical areas using MRS before, during and after learning in children and adults for the first time, we found that an initially lower concentration of GABA in children than in adults rapidly boosted only in children after learning. This rapid boosting of GABA in children leads to the prediction that children more rapidly stabilize learning than adults, which is indeed supported by subsequent behavioral experiments.

In the current study, we found the GABA boosting effect after learning in school age children. It is known that GABA plays a crucial role early in development during the visual critical periods (*28–30*). It would be interesting to examine whether a similar GABA boosting effect also occurs in infants during the visual critical periods and if so, whether and how the effect influences the visual critical periods.

## Acknowledgments

We wish to thank the children and their parents for participating in this study.

## Authors contributions

Conceptualization: SMF, YS, MWG, TW

Methodology: SMF, WMM

Investigation: SMF, MB, AQ, PG, UIF, WMM

Visualization: SMF

Funding acquisition: TW, MWG

Project administration: SMF, YS, MWG, TW

Supervision: YS, MWG, TW

Writing – original draft: SMF, TW

Writing – review & editing: SMF, YS, MWG, TW

## Data and materials availability

The original data are available online.

## Supplementary Materials

Materials and Methods

Figs. S1-8

References 31–44

## Supplementary Materials

## Materials and Methods

### Subjects

Taken together over all experiments, a total of 112 subjects (56 children and 56 adults) were recruited for this study. The results of one child in the MRS experiment could not be used due to MRS measurement failure. The remaining subjects in the group of children (30 female and 25 male, 53 right-handed) had a mean ± SEM age of 9.64 ± 0.14 years (range: 8 – 11 years old). Adult subjects (36 female and 20 male, 49 right-handed) had a mean ± SEM age of 22.5 ± 0.61 years (range: 18 – 35 years old). For each experiment (i.e., preliminary experiment to determine sample size, MRS experiment, behavioral experiments with 10 min and 60 min intermissions) a total of 14 subjects per age group were recruited. The number of subjects was determined by means of a power analysis (see *Preliminary Experiment to Determine Sample Size*). All subjects had normal or corrected-to-normal vision. Visual acuity in children was determined as normal in a state-regulated medical screening required to enroll in elementary school. The study was approved by the local ethics committee. Informed written consent was obtained from each subject. For the group of children informed written consent was additionally provided by their legal guardian.

### Training Task

A two-interval-forced-choice orientation detection task (Fig. 1A) was used for training in each experiment, following the approach of previous studies (*2, 3*). Each trial started with a central fixation spot (diameter: 0.675°) presented for 500 ms. Thereafter, two different images were flashed for 50 ms each, separated by a blank interval of 300 ms. One image consisted of a Gabor pattern presented at the center of the screen with a specific orientation overlaid by a noise pattern generated from a sinusoidal luminance distribution. The noise pattern replaced a subset of pixels in the original Gabor pattern. The Gabor pattern had the following parameters: contrast = 100% (rounded), spatial frequency = 1 cycle per degree, sigma of Gaussian filter = 2.5°, random spatial phase, diameter = 4.5°. The orientation of the Gabor pattern was either 10°, 70° or 130° away from the vertical orientation. The other image consisted of a pure noise pattern of the same size as the Gabor pattern and was also presented at screen center. In each trial the two different images were presented in random order. Subjects were asked to maintain central fixation and to indicate at trial end which of the two images contained the orientation by pressing one of two buttons on the keyboard during the behavioral experiments and the button-box during the MRS experiment. No feedback about response accuracy was provided. Task difficulty was controlled by a one-up-two-down adaptive staircase procedure yielding an average response accuracy of 70.7% correct. Task difficulty was adjusted by varying the subset of pixels in the Gabor pattern replaced by noise. The more pixels replaced by noise the more difficult was the task. In the first trial of each block of the task 25 percent of the original pixels in the Gabor pattern were replaced by noise. This ratio was adjusted with a step size of 0.05-log units during the adaptive staircase. Around 40 – 50 trials (terminated after ten reversals) were conducted for each block of the task (taking between 2 – 3 min). The orientation detection threshold for each block was calculated as the geometric mean of the last six reversals.

Stimuli were generated using Psychtoolbox (*31, 32*) running in MATLAB (The Mathworks, Nattick, MA, USA) and presented using a gamma-corrected LCD screen in a behavioral testing room with lights turned off. Subjects were seated 60 cm away from the screen and used a chin rest. For the MRS experiment stimuli were projected using a projector with gamma-correction onto a translucent screen located at the back of the scanner bore. Subjects viewed the screen with a headcoil-mounted mirror.

### Preliminary Experiment to Determine Sample Size

A preliminary experiment was conducted in children and adults to determine the sample size for the experiments in this study. The number of subjects for the preliminary experiment was determined based on a previous study (*2*), which employed a similar design and training task as in the preliminary experiment. In this original study a total of 12 subjects were recruited. To be a little more conservative, we decided to run a total of 14 subjects for each age group in the preliminary experiment. Each subject completed two sessions on separate days. The first session consisted of pretest and training. For pretest, subjects completed three successive blocks of the orientation detection task using the same orientation in each block (in the following referred to as the trained orientation). The trained orientation was randomly chosen from the set of three different orientations (see above) for each subject.

The first block of pretest served as practice and the respective results were not further analyzed (following the approach by *3*). Immediately following pretest, subjects performed eight successive blocks of the orientation detection task using the trained orientation. The second session consisted of a posttest with the same design as the pretest. VPL was calculated as the percent improvement in detection threshold for the trained orientation from pretest to posttest using the following formula: (1 – threshold at posttest / threshold at pretest) x 100. The thresholds in pretest and posttest corresponded to the arithmetic mean across thresholds in the second and third blocks of each test. If subjects exhibited VPL, their detection thresholds for the trained orientation should improve from pretest to posttest. The results showed that children and adults both exhibited a significant improvement of detection thresholds for the trained orientation from pretest to posttest. The mean ± SEM percent threshold improvement from pretest to posttest was 19.8 ± 5.22% across children and 19.8 ± 5.70% across adults. One-sample *t*-tests of percent threshold change in posttest against zero corresponding to pretest threshold showed a significant improvement of thresholds in children [*t*(13) = 3.79, *p* = 0.002, *d* = 1.01] and adults [*t*(13) = 3.48, *p* = 0.004, *d* = 0.93], indicative of VPL (see *2, 3*, for similar results in adults). The magnitude of improvement (i.e., percent threshold change from pretest to posttest) was not significantly different between children and adults [two-sample *t*-test; *t*(26) = −0.01, *p* = 0.995].

The changes in thresholds from pretest to posttest for each age group were submitted to a power-analysis to determine the sample size for the experiments in this study. We calculated the number of subjects necessary to achieve a power of 0.80 at alpha = 0.05 (two-tailed) using a one-sample *t*-test to detect the minimum effect size of interest for VPL of the trained orientation in each age group. The results of the power analysis (using G*Power; *33*) showed that a total of 10 subjects were necessary to detect a minimum effect size of interest (*d* = 1.01, calculated from the preliminary experiment) for VPL of the trained orientation in children. A total of 12 subjects were necessary to detect a minimum effect size of interest (*d* = 0.93, calculated from the preliminary experiment) for VPL of the trained orientation in adults. Based on these results, we decided to follow the approach of the preliminary experiment and to recruit a total of 14 subjects for each age group in each experiment.

### MRS Experiment

The experiment consisted of MRS measurements prior to (pretraining MRS), during (training MRS), and immediately after (posttraining MRS) training on a VPL task. MRS was conducted for a voxel located in early visual cortical areas (Fig. 1C). Pretraining MRS served as baseline to measure the concentration of GABA prior to VPL. Training MRS was conducted while subjects trained on the VPL task. Posttraining MRS was conducted immediately after the end of training on the VPL task.

MRS was conducted in scans of ∼ 3 min each (see *Imaging Parameters* below for details). A total of 13 successive MRS scans were conducted for each subject. Three successive scans were conducted for pretraining MRS [corresponding to 0 – 3 min (MRS1), 3 – 6 min (MRS2) and 6 – 9 min (MRS3), relative to the onset of pretraining MRS], resulting in a total pretraining scan time of ∼ 10 min. Four successive MRS scans were conducted for training MRS [corresponding to 0 – 3 min (MRS4), 3 – 6 min (MRS5), 6 – 9 min (MRS6) and 9 – 12 min (MRS7), relative to the onset of training MRS], resulting in a total training scan time of ∼ 13 min. Six successive scans were conducted for posttraining MRS [corresponding to 0 – 3 min (MRS8), 3 – 6 min (MRS9), 6 – 9 min (MRS10), 9 – 12 min (MRS11), 12 – 15 min (MRS12) and 15 – 18 min (MRS13), relative to the onset of posttraining MRS], resulting in a total posttraining scan time of ∼ 20 min. All MRS measurements were conducted successively without any intermissions. In pretraining and posttraining MRS subjects performed a speeded color change detection task at central fixation to maintain subjects’ fixation and attention at screen center during the MRS scans, similar to the approach of previous studies (*2, 3*). To this purpose a white dot (diameter = 3°) placed inside a gray disk was presented at screen center. Occasionally, in an unpredictable fashion, the white disk changed color briefly (for 1.5 s) from white (R, G, B = 255, 255, 255) to faint pink (R, G, B = 255, 255-X, 255-X) and then returned to white (X refers to the G and B intensity change, see below). Subjects were asked to press a button on the buttonbox whenever they detected a color change from white. If subjects pressed a button within the 1.5 s long period of a color change, the response was considered correct. If subjects did not press a button within this period or if they pressed a button after this period, this was considered an incorrect response. The task difficulty was controlled by using a one-up-two-down adaptive staircase procedure. The initial color change X was set to 40. The color change was adjusted by either decreasing or increasing the magnitude of color change in a step size of 2 to increase or decrease task difficulty, respectively. Task performance was analyzed by calculating the color change detection threshold for each scan of pretraining and posttraining MRS. This threshold corresponded to the geometric mean of the color change values across the last six reversals for each scan.

Figure S1 shows the mean (± SEM) thresholds to detect a color change of the central fixation spot for each scan of pretraining and posttraining MRS. Mann-Whitney-U tests did not show any significant differences in thresholds between children and adults in any MRS scan [Pretraining 0 – 3 min: *U* = 48.5, *p* = 0.35; Pretraining 3 – 6 min: *U* = 74.5, *p* = 0.61; Pretraining 6 – 9 min: *U* = 71.5, *p* = 0.61; Posttraining 0 – 3 min: *U* = 59.0, *p* = 0.47; Posttraining 3 – 6 min: *U* = 62.0, *p* = 0.47; Posttraining 6 – 9 min: *U* = 78.5, *p* = 0.61; Posttraining 9 – 12 min: *U* = 68.5, *p* = 0.61; Posttraining 12 – 15 min: *U* = 89.5, *p* = 0.94; Posttraining 15 – 18 min: *U* = 71.5, *p* = 0.61; all *p*-values after FDR-correction]. These results show that children and adults exhibited similar performance in the fixation task during pretraining and posttraining MRS, indicating that both age groups fixated and attended equally well at screen center during scanning. Therefore, we deem it unlikely that differences in GABA between children and adults were influenced by differences in fixation quality or attention during the MRS measurements between the age groups (see also *2, 3*, for similar results using the same color change detection task during MRS).

**Fig. S1.**
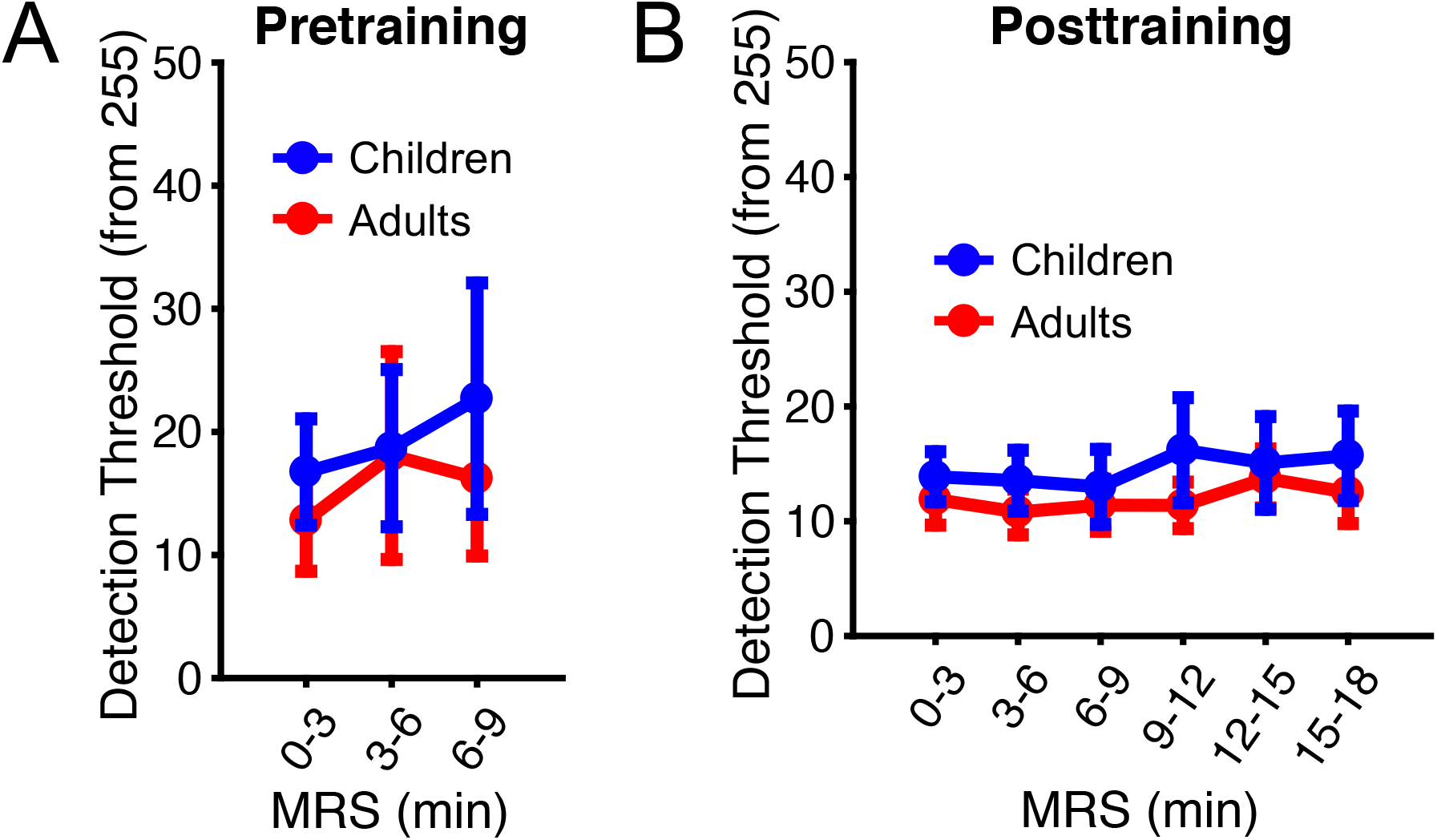
Performance in the color change detection task. (**A**) Mean (± SEM) thresholds to detect a color change of the central fixation spot across children (n = 13) and adults (n = 14) during pretraining. MRS scans were collected every 3 min. Three scans were conducted for pretraining MRS. 0 min on the x-axis corresponds to the onset of the first pretraining scan. The y-axis shows the threshold in units of change of green and blue intensities from 255. (**B**) Same as (A) but for posttraining. Six scans were conducted for posttraining. 0 min on the x-axis corresponds to the onset of the first posttraining scan.

The same orientation detection task as in the preliminary experiment served as the VPL task for the MRS experiment. Each subject was tested and trained on an orientation that was randomly chosen from the set of three orientations for each subject. One block was conducted for pretest. Eight blocks were conducted for training. On a separate day a posttest in the trained task was conducted exactly as the pretest but outside the scanner. VPL was calculated as in the preliminary experiment (see *Preliminary Experiment to Determine Sample Size*). Pretest detection thresholds for the trained orientation were not significantly different between children and adults (Mann-Whitney U-test; *U* = 85.0, *p* = 0.77).

### Imaging Parameters

Imaging was conducted using a Siemens 3T Prisma MRI scanner (Siemens, Erlangen, Germany) using a 64-channel head/neck coil. High-resolution anatomical MRI scans of the brain were collected using three sequences for coronal, sagittal and transverse planes to support the placement of the MRS voxel (see below). These sequences had the following imaging parameters: coronal plane [time-to-repeat (TR) = 0.25 s, time-to-echo (TE) = 2.46 ms, flip angle (FA) = 70°, in-plane acquisition matrix (AM) = 288 × 288, 35 slices, voxel size = 0.8 × 0.8 × 4.0 mm, inter-slice gap = 1.20 mm], sagittal plane (TR = 0.19 s, TE = 2.46 ms, FA = 70°, AM = 288 × 288, 25 slices, voxel size = 0.8 × 0.8 × 4.0 mm, inter-slice gap = 1.20 mm), transverse plane (TR = 0.19 s, TE = 2.46 ms, FA = 70°, AM = 288 × 288, 27 slices, voxel size = 0.8 × 0.8 × 4.0 mm, inter-slice gap = 1.20 mm). The sagittal scan was used to reconstruct and inflate each subject’s brain using the Freesurfer software package (Martinos Center for Biomedical Imaging, Charlestown, MA, USA; *34, 35*). Freesurfer’s segmentation of the brain into gray matter (GM) and white matter (WM) during reconstruction was used for subsequent volume fraction calculations and corrections of the estimated metabolite concentrations (see *MRS Analysis*). The anatomical segmentation of the cerebral cortex proposed by Glasser and colleagues (*36*) was remapped from the Freesurfer average brain onto each subject’s reconstructed brain to identify cortical areas included in the GM of the MRS voxel.

Single-voxel proton (^1^H) MRS was conducted for a voxel (2 × 2 × 2 cm) placed manually alongside the calcarine sulcus. The voxel was oriented perpendicular to the calcarine sulcus and centered between the two hemispheres (Fig. 1C). MRS scans were collected using a MEGA-PRESS sequence (*37, 38*) with the following imaging parameters: TR = 1.5 s; TE = 68 ms; FA = 90°; number of averages = 64; scan time = 3:18 min. The number of averages followed the approach of a previous study (*25*). A frequency selective, single band Gauss pulse was utilized to saturate the *β*-CH2 signal at 1.94 ppm and to refocus the J evolution of the triplet γ-CH2 resonance of GABA at 3 ppm (‘Edit On’). The very same Gauss pulse was used to irradiate the opposite part of the spectrum at 7.46 ppm (‘Edit Off’). The ‘Edit Off’ spectrum was subtracted from the ‘Edit On’ spectrum to produce a difference spectrum. The concentrations of GABA and the composite measure of N-acetyl-aspartate and N-acetyl-aspartyl-glutamate (NAA+NAAG) were estimated from this difference spectrum (see below). WET water suppression was used (*39*). Furthermore, a water reference scan for the same voxel was collected using a PRESS sequence without water suppression (TR = 3 s; TE = 30 ms; FA = 90°; number of averages = 16; scan time = 1:03 min).

Non-neural tissue containing lipids was avoided during voxel placement. For each subject volume fractions inside the voxel corresponding to three categories (GM, WM, others) were calculated. The mean volume fractions (± SEM) of GM, WM inside the voxel across subjects in each age group of the MRS experiment were (fig. S2a): GM = 73.1 ± 2.66% (children), 67.3 ± 3.10% (adults); WM = 26.9 ± 2.65% (children) and 32.0 ± 3.08% (adults); Other types of volume were negligible (see fig. S2a). Two-sample *t*-tests showed no significant differences in the amount of GM [*t*(25) = 1.40, *p* = 0.17] and WM [*t*(25) = −1.25, *p* = 0.22] within the voxel between children and adults. The GM of the voxel corresponded primarily to early visual area V1 and to a lesser degree area V2 (fig. S2b) as defined by the segmentation of Glasser and colleagues (*36*). GM corresponding to V3 (again, using the segmentation by *36*) was negligible in each age group. Two-sample *t*-tests showed no significant differences in the amount of V1 [*t*(25) = −0.72, *p* = 0.47] and V2 [*t*(25) = 0.57, *p* = 0.57] within the GM of the voxel between children and adults.

**Fig. S2.**
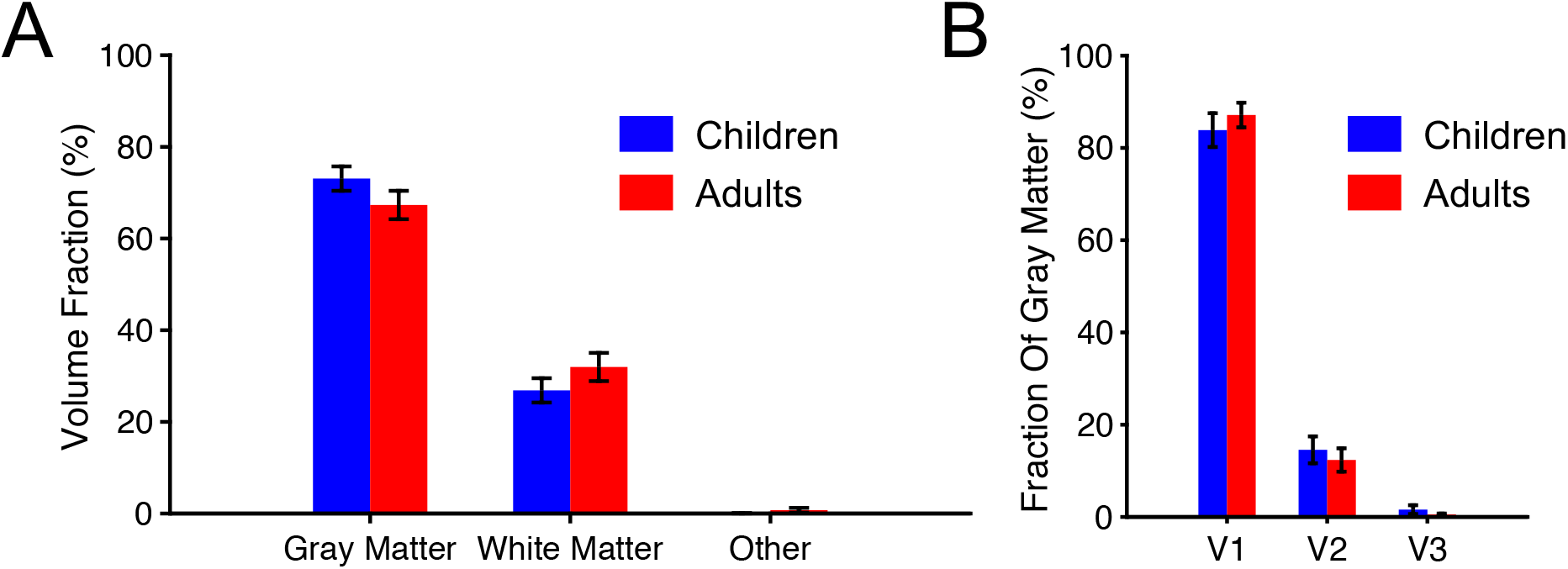
Volume fractions and early visual cortical areas within the voxel used for MRS. (**A**) Mean (± SEM) fractions of gray matter, white matter and other volume types within the voxel across children (n = 13) and adults (n = 14) in percent. (**B**) Early visual cortical areas within the gray matter of the voxel. Otherwise same as (A).

Automatic shimming, i.e. gradient field adjustments required to increase the homogeneity of the magnet field B0, was conducted prior to each MRS scan. If necessary, manual shimming was conducted after automatic shimming. We aimed to keep the shimming values for each subject and MRS acquisition below a full-width at half-maximum (FWHM) of 20 Hz. This was possible in all but six scans (out of 243 scans collected in total across all subjects). The shimming values in these six scans ranged between 20.1 to 25.6 Hz. The mean shim values (with SEM) for each scan across subjects in each age group are shown in fig. S3. For one subject from the group of children the last scan could not be conducted and hence there is no shim value for this scan in this subject. A 13 × 2 mixed design ANOVA with the within-subject factor of MRS scan (MRS1 – MRS13) and the between-subject factor of age group (children, adults) did not show any significant main effects of MRS scan [*F*(12,288) = 1.53, epsilon = 0.32, *p* = 0.20] or age group [*F*(1,24) = 2.60, *p* = 0.12]. Furthermore, there was no significant interaction between MRS scan and age group [*F*(12,288) = 1.25, epsilon = 0.32, *p* = 0.30]. These results indicate that shim values in the MRS experiment were similar across scans and between the age groups.

**Fig. S3.**
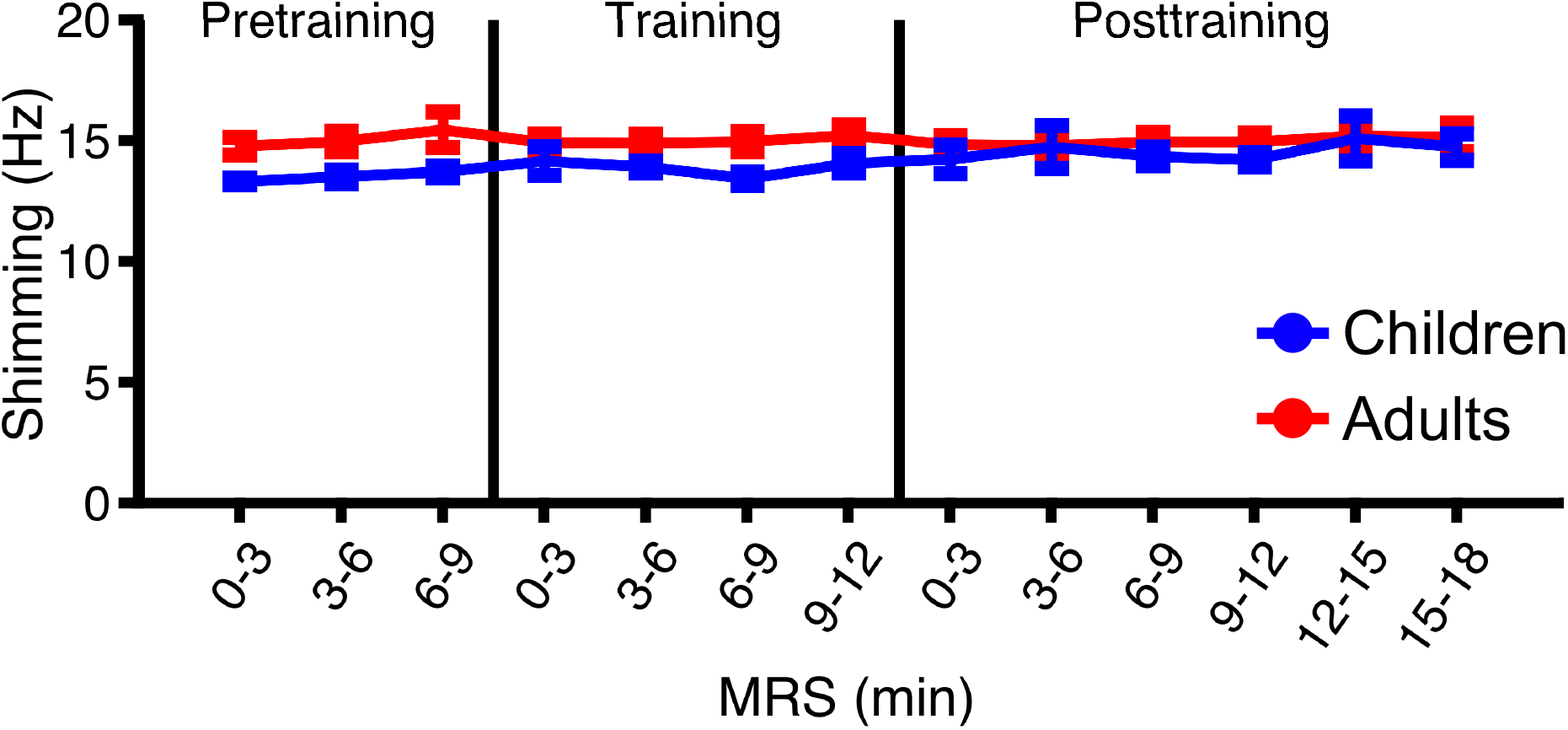
Shimming results. Mean (± SEM) shim values (in Hz) for each MRS scan across children (n = 13) and adults (n = 14). MRS scans were collected every ∼ 3 min. Three scans were collected for pretraining. 0 min on the x-axis in pretraining corresponds to the onset of the first pretraining scan. Four scans were collected for training. 0 min on the x-axis in training corresponds to the onset of the first training scan. Six scans were collected for posttraining. 0 min on the x-axis in posttraining corresponds to the onset of the first posttraining scan.

### MRS Analysis

The metabolite concentrations of GABA and NAA+NAAG measured by MEGA-PRESS were analyzed with the LC model (*40, 41*) in the chemical shift range between 1.9 and 4.2 ppm. NAA is the hydrolysis product of NAAG and a marker of neuronal density and mitochondrial function (*42*). It is commonly used as a control metabolite to normalize the concentration of GABA (*18, 19*). Figures S4 and S5 show the individual MEGA-PRESS difference spectra for each MRS scan and subject of each age group. Water scaling and eddy current correction were performed. The metabolite intensities were fitted to a linear combination of spectra of individual metabolites derived from an imported metabolite basis set (*40, 41*). The Cramer-Rao Lower bounds (CRLB) indicate the reliability of quantification of GABA and NAA+NAAG. We used a criterion of 25% to reject low-quality results. Using this criterion, a total of 13 scans for GABA across subjects from the group of children [mean CRLB% (± SEM) across excluded scans: 30.5% (± 1.56)] and a total of 10 scans for GABA across subjects from the group of adults [mean CRLB% (± SEM) across excluded scans: 31.2% (± 1.58)] had to be excluded. The percentage of excluded scans relative to all scans collected across each age group was: 7.73% (group of children) and 5.49% (group of adults). None of the results had to be excluded for NAA+NAAG in any subject. The mean CRLB% (± SEM) across the remaining scans and subjects in each age group were 18 ± 0.37% (children) and 18 ± 0.34% (adults) for GABA and 2 ± 0.04% (children) and 2 ± 0.05% (adults) for NAA+NAAG (see fig. S6 for mean values for each scan across subjects in each age group).

The absolute concentrations of GABA and NAA+NAAG from the difference spectrum in each scan were corrected for each subject’s volume fractions inside the MRS voxel (following the approach by *43, 44*). Specifically, GABA was corrected for the proportion of GM within the voxel by dividing by [GM / (GM + WM + Other)]. The concentration of NAA+NAAG was corrected for the proportion of total brain volume within the voxel by dividing by [(GM + WM) / (GM + WM + Other)]. Then, for each subject and scan the corrected absolute concentration of GABA was normalized to the corrected absolute concentration of NAA+NAAG. NAA+NAAG was highly stable across MRS scans in each age group as shown by a repeated measures ANOVA with the factor of MRS scan (MRS1 – MRS13) on the corrected absolute concentration of NAA+NAAG in the MRS experiment. There were no significant differences in NAA+NAAG across MRS scans in children [*F*(12,132) = 0.79, *p* = 0.67] and adults [*F*(12,156) = 1.41, *p* = 0.16].

The corrected and normalized metabolite concentrations of GABA were averaged across pretraining MRS1 – MRS3 for each subject and used as baseline to calculate changes in the concentration of GABA from pretraining to posttraining in percent. Changes from pretraining to posttraining were calculated across posttraining MRS (corresponding to the corrected and normalized metabolite concentrations averaged across posttraining MRS8 – MRS13 for each subject; Fig. 1D) and separately for each scan of posttraining MRS (Fig. 1E). If a scan could not be used in a given subject either because of a CRLB exceeding our inclusion criterion (see above) or because of a collection failure (for one child subject), the averaging was performed across the remaining scans. See fig. S7 for mean corrected and normalized concentrations of GABA for each scan across subjects in each age group.

**Fig. S4.**
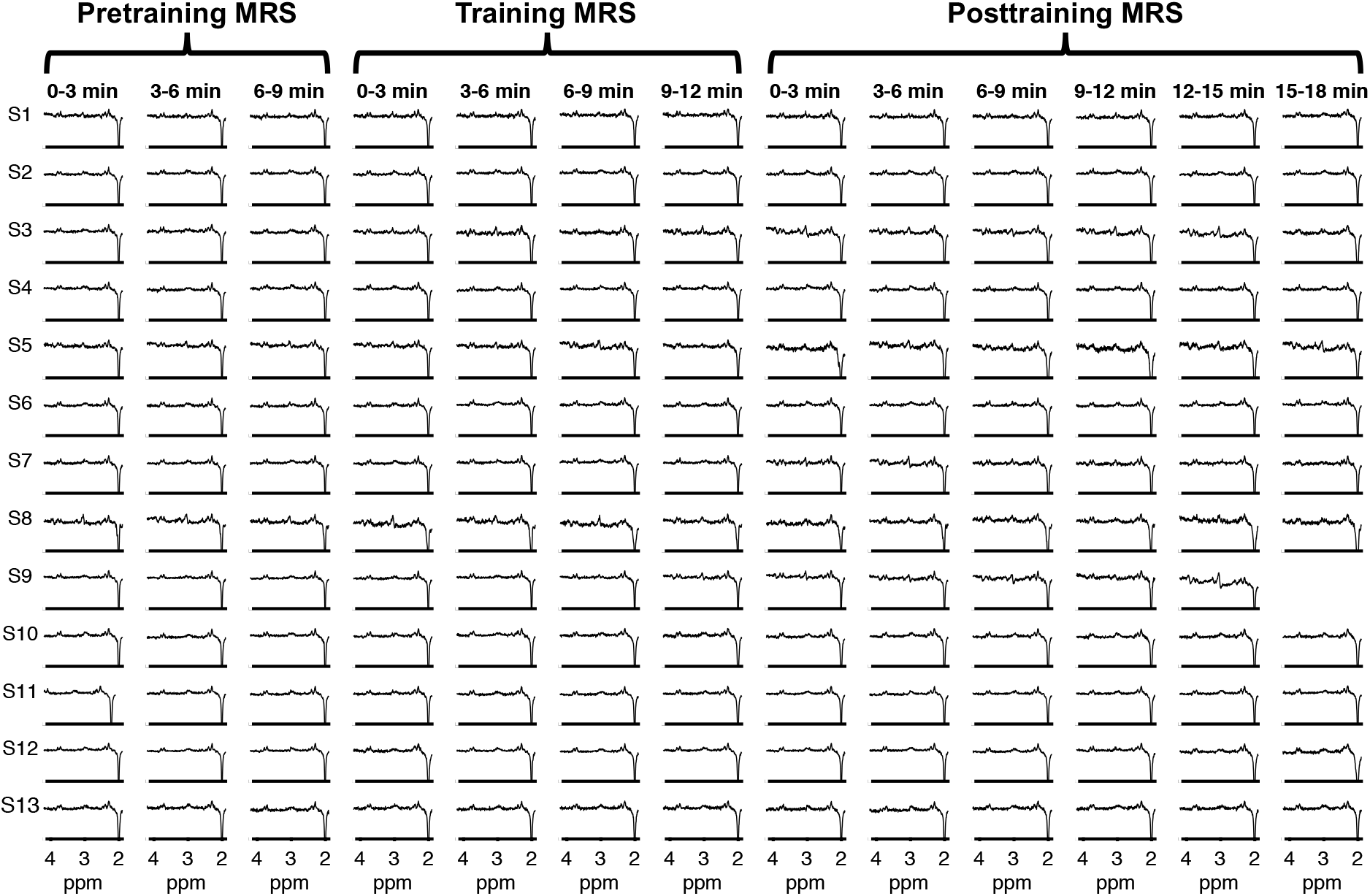
MEGA-PRESS difference spectrum for each MRS scan and subject in the group of children. Time-points of pretraining, training and posttraining MRS as in fig. S3. Each row shows the result of a different child subject (S1 – S13). The 15 – 18 min posttraining scan could not be collected for S9. The GABA peak is seen at ∼ 3ppm. The NAA+NAAG peak is seen at ∼ 2ppm.

**Fig. S5.**
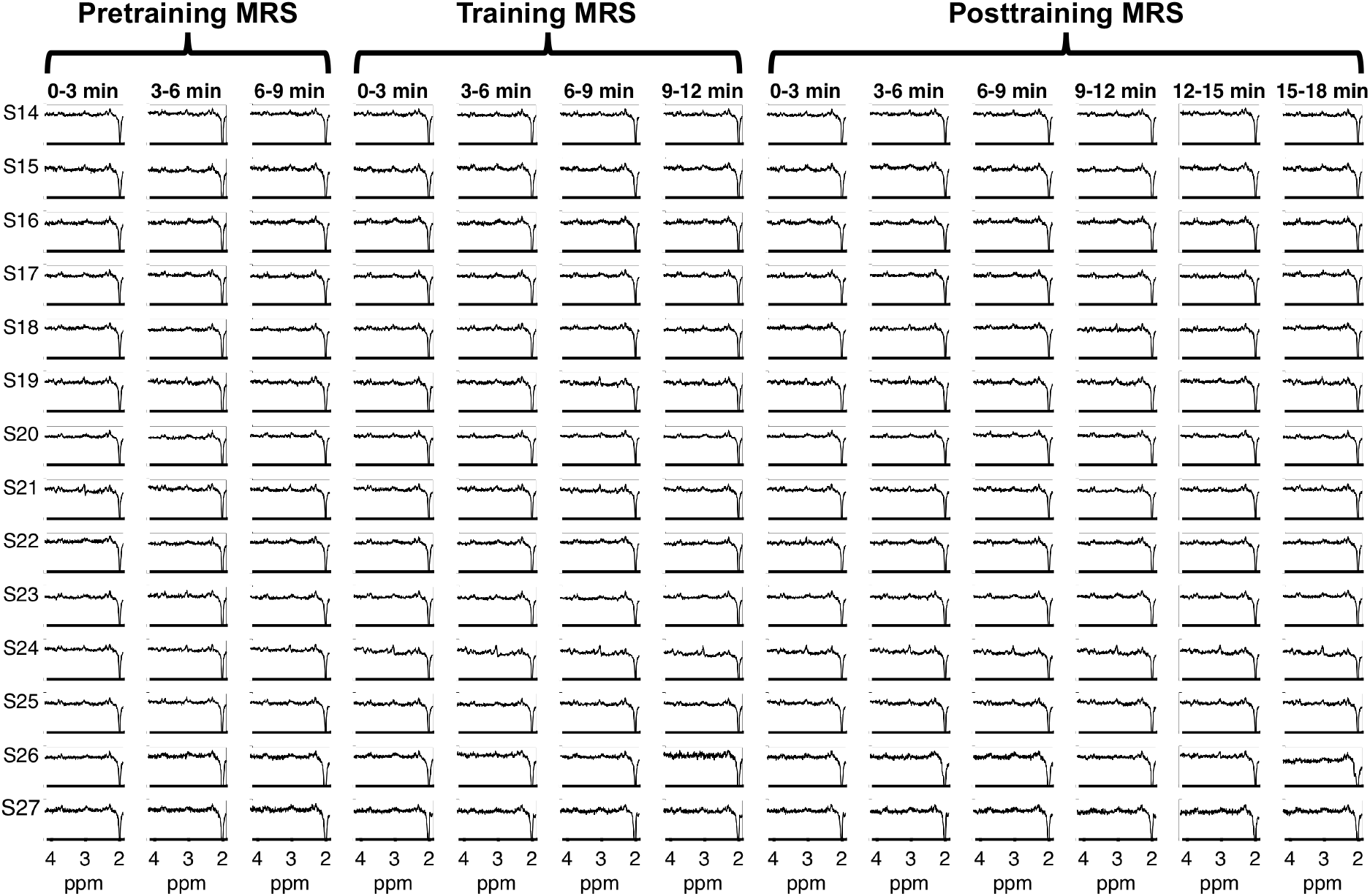
Same as fig. S4 but for adult subjects (S14 – S27).

**Fig. S6.**
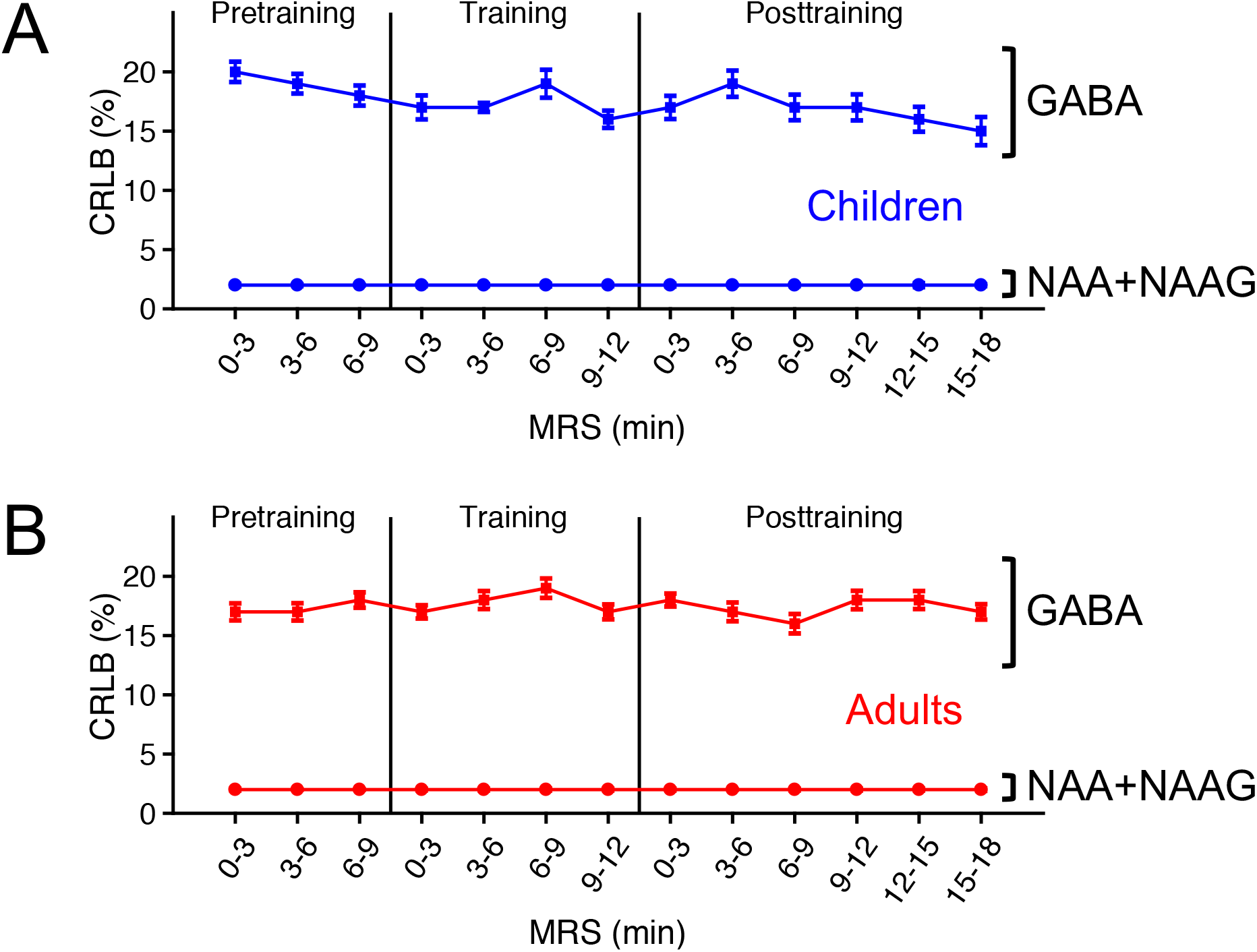
LC model fitting results. (**A**) Mean (± SEM) uncertainties [corresponding to Cramer-Rao Lower bounds (CRLB) in percent] of different metabolites (GABA, NAA+NAAG) for each MRS scan across children (n = 13). Time-points of pretraining, training and posttraining MRS as in fig. S3. SEMs of CRLBs for NAA+NAAG were close to zero. (**B**) Same as (A) but for the group of adults (n = 14).

**Fig. S7.**
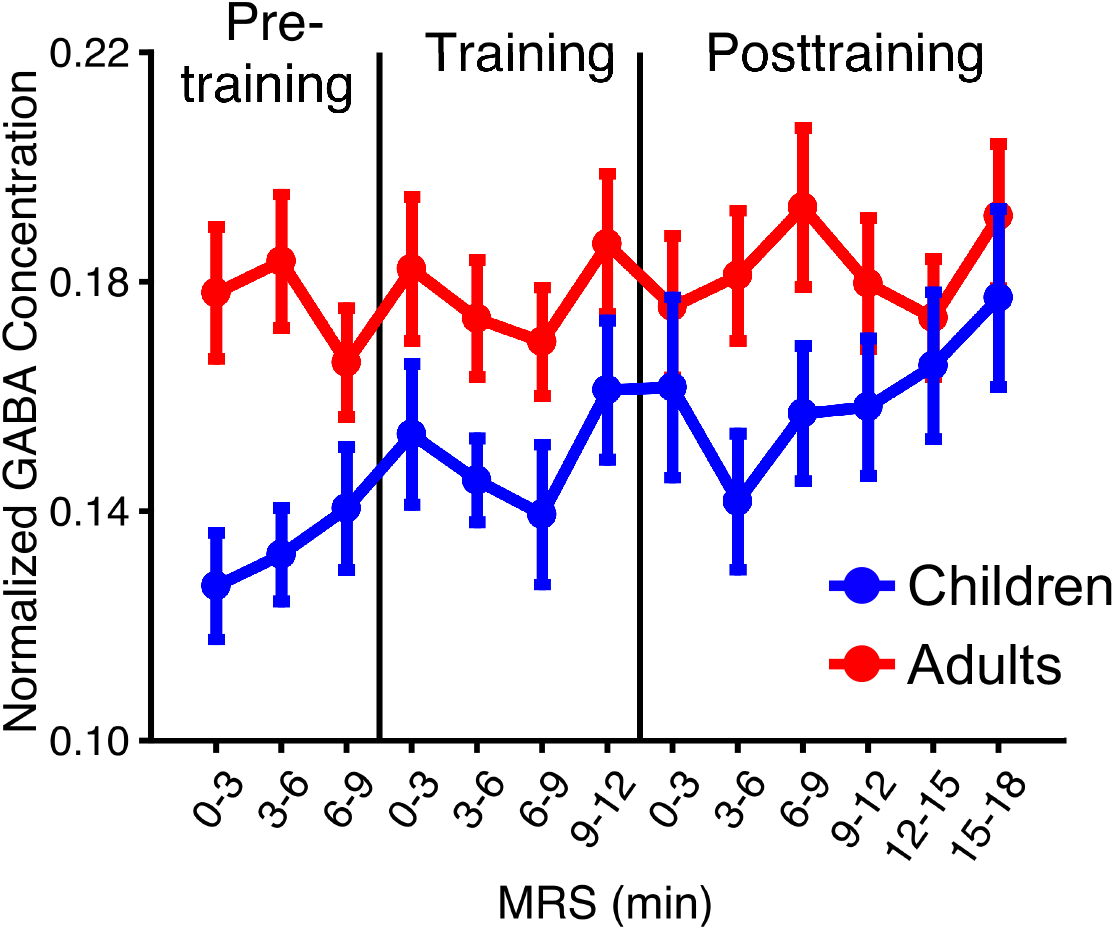
Normalized GABA concentrations. Mean (± SEM) concentrations of GABA normalized to NAA+NAAG for each MRS scan across children (n = 13) and adults (n = 14). Time-points of pretraining, training and posttraining MRS as in fig. S3.

### Behavioral Experiments with 10 min and 60 min Intermissions

We conducted two behavioral experiments, which only differed in the duration of the intermission between training of the first and second trained orientations (see below). Each experiment consisted of two sessions on separate days (Fig. 2A and fig. S8a). The same orientation detection task as in the preliminary experiment was used. Subjects were tested and trained on two different orientations, which were randomly chosen from the set of three different orientations for each subject. In the first session subjects performed a pretest for each of the two trained orientations. Three successive test blocks were conducted for each trained orientation and the order in which the two orientations were tested was random for each subject. As in the preliminary experiment the first test block for each trained orientation served as practice and was omitted from further analyses. Pretest thresholds were calculated as the arithmetic mean across thresholds in the second and third test blocks. Immediately following pretest subjects trained for eight successive blocks on the first trained orientation.

After the end of this training, intermissions of 10 min and 60 min were included in different experiments. These durations were chosen, because previous studies showed that adults exhibited retrograde interference in VPL when they trained on a second VPL task within an interval of up to one hour after training on a first VPL task (*2, 13*). Therefore, we predicted that adult subjects would exhibit retrograde interference with intermissions of both 10 min and 60 min in the current experiments. During the intermission subjects did not perform any training and were free to leave the behavioral testing room. Thereafter, subjects performed another eight successive training blocks on the second trained orientation. In the second session subjects performed a posttest for each trained orientation exactly as during pretest. As in the pretest, the first block of posttest served as practice and was not any further analyzed. Posttest thresholds were calculated as the arithmetic mean across thresholds in the second and third test blocks. The order in which each trained orientation was tested during posttest was random for each subject. Changes in thresholds to detect each trained orientation from pretest to posttest were calculated exactly as in the preliminary experiment.

The results of the experiment with a 10 min intermission are shown in Fig. 2B. The results of the experiment with a 60 min intermission showed that children improved detection thresholds for each trained orientation from pretest to posttest [one-sample *t*-test of percent threshold change in posttest against zero corresponding to pretest threshold; first trained orientation: *t*(13) = 3.91, *p* = 0.002, *d* = 1.05; second trained orientation: *t*(13) = 3.11, *p* = 0.008, *d* = 0.83] (fig. S8b). In contrast to children, adults improved detection thresholds from pretest to posttest only for the second trained orientation [*t*(13) = 2.56, *p* = 0.02, *d* = 0.68] but not for the first trained orientation [*t*(13) = 0.83, *p* = 0.42] (fig. S8b). This difference in percent threshold change from pretest to posttest for the first trained orientation between children and adults was significant [two-sample *t*-test; *t*(26) = 2.24, *p* = 0.03, *d* = 0.85], whereas no significant difference between the age groups was found for the second trained orientation [*t*(26) = 1.42, *p* = 0.17]. There were no significant differences in pretest detection thresholds between children and adults for the first [*t*(26) = 1.69, *p* = 0.10] and second trained orientations [*t*(26) = 1.00, *p* = 0.33].

**Fig. S8.**
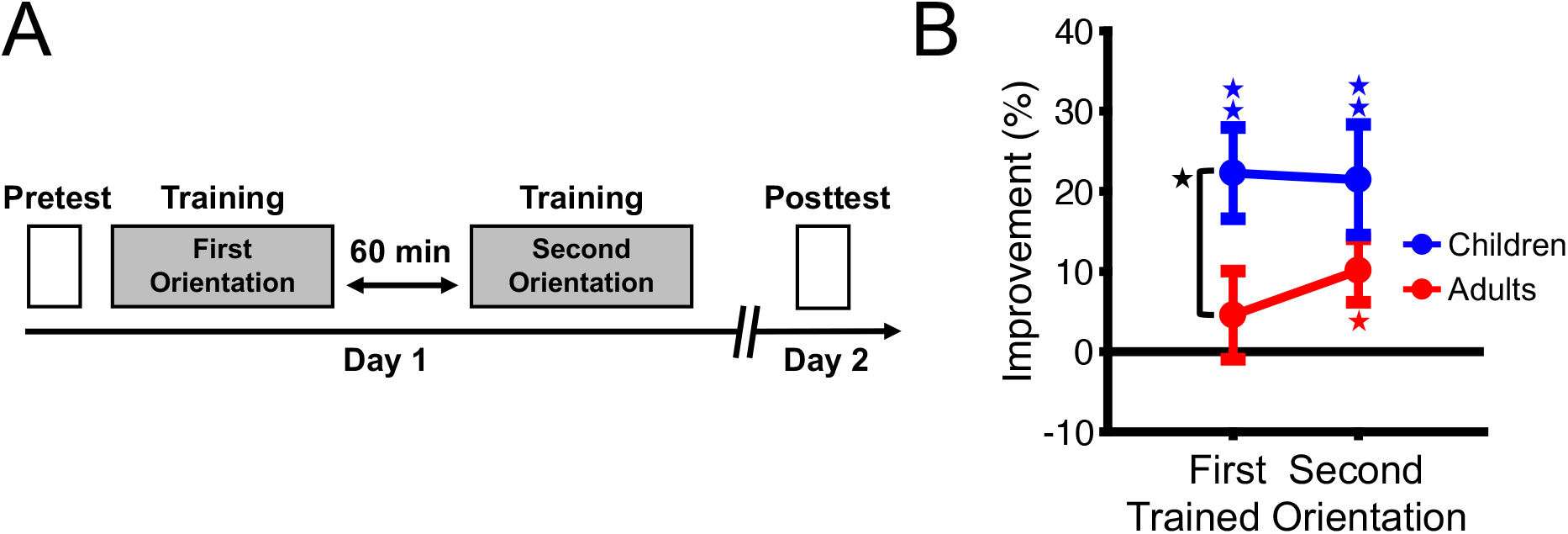
Design and results of the behavioral experiment with 60 min intermission between trainings. (**A**) Design of the experiment. (**B**) Results of the experiment. Mean (± SEM) changes in detection thresholds from pretest to posttest for each trained orientation across children (n = 14) and adults (n = 14). Zero on the y-axis represents pretest thresholds. Values above zero indicate that thresholds improved from pretest to posttest. Blue and red asterisks indicate a significant improvement in thresholds from pretest to posttest, indicative of visual perceptual learning, for children and adults, respectively. The black asterisk between children and adults for the first trained orientation indicates that improvements for this orientation were significantly larger in children than adults. * *p* < 0.05, ** *p* < 0.01.

### Statistics

The results of this study were analyzed using parametric statistics, after confirmation of the assumption of normality using the Shapiro-Wilk test. The assumption of normality was violated for pretest thresholds in the MRS experiment and for thresholds in the color change detection task, as shown by significant results of the Shapiro-Wilk test. Therefore, non-parametric Mann-Whitney-U tests were used for the statistical analysis of these results. If the assumption of sphericity for ANOVA was violated as shown by a significant result in Mauchly’s test of sphericity, the Huynh-Feldt correction was used. The effect sizes of significant results are reported as partial *η*^2^ for ANOVA and Cohen’s *d* for *t*-test. For all statistical tests the two-tailed alpha level was set to 0.05.

